# «Application of RT-LAMP-CRISPR-Cas13a technology to the detection of OXA-48 producing *Klebsiella pneumoniae*»

**DOI:** 10.1101/2022.08.29.505698

**Authors:** Concha Ortiz-Cartagena, Lucia Blasco, Laura Fernández-García, Olga Pacios, Ines Bleriot, María López, Felipe Fernández-Cuenca, Rafael Cantón, María Tomás

## Abstract

Carbapenem-resistant pathogens have been recognized as a health concern because of their ability to cause severe infections and because they are difficult to detect in laboratories. Researchers are making great efforts to develop a diagnostic technique with high levels of sensitivity and specificity, as accurate, early diagnosis is required to prevent the spread of these microorganisms and improve the prognosis of patients. In this context, CRISPR-Cas systems are proposed as promising tools for the development of diagnostic techniques due to their high specificity: Cas13 endonuclease discriminates single nucleotide changes and displays collateral activity against single-stranded RNA molecules. This technology is usually combined with isothermal pre-amplification reactions in order to increase the sensitivity of diagnosis. We have developed an RT-LAMP-CRISPR-Cas13a-based assay for the detection of *Klebsiella pneumoniae* OXA-48 producer strains in clinical samples without the need for RNA extraction. The assay exhibited 100 % specificity, sensitivity, positive predictive value and negative predictive value.

## Introduction

Bacterial antimicrobial resistance (AMR) has emerged as a threat to human public health in the 21^st^ century, being responsible for 1,27 million deaths in 2019 due to the severe infections it causes in immunocompromised patients [1, 2]. Furthermore, these illnesses lead to long-term hospitalization of patients, involving expensive and intensive care [3, 4]. The ESKAPE bacteria (*Enterococcus faecium, Staphylococcus aureus, Klebsiella pneumoniae, Acinetobacter baumannii, Pseudomonas aeruginosa* and *Enterobacter* spp.) stand out due to the enhanced capacity of the members of this group to gain plasmids with antibiotic-resistant genes. The ESKAPE group was responsible for 929,000 deaths in 2019 [1].

Carbapenem antibiotics have become drugs of last resort (DoLR) to treat severe infections caused by Gram-negative bacteria [5-8]. Worryingly, reports of Carbapenem-Resistant Enterobacterales (CRE), which are included in the list of priority pathogens compiled by the World Health Organization (WHO) [9], are constantly increasing. Among the CRE, *K. pneumoniae* most frequently shows resistance to these antibiotics, causing between 50,000 and 100,000 deaths in 2019 [1]. The Enterobacterales have developed three different carbapenem resistance mechanisms, being the expression of carbapenemase enzymes the principal [3, 8, 10]. The most important carbapenemases in the Enterobacteriaceae are *K. pneumoniae* carbapenemase (KPC)-like enzymes (Class A), metallo-β-lactamases (Class B) (VIM, IMP, NDM) and OXA carbapenemases (Class D) [3, 10]. OXA-48-like lactamases (Class D) are very common among global Enterobacterales, and in some areas, such as Spain, OXA-48-like enzymes are the primary carbapenemases in this order and are mainly present in mobile genetic elements of *K. pneumoniae* [8, 11]. Besides, carbapenem-resistant *K. pneumoniae* (CRKP) has been recovered from hospital environments and also from community samples (drinking water and food-producing animals), and hypervirulent *Klebsiella* strains with carbapenemase enzyme-encoding genes have also been detected. These findings both increase the spectrum of people at risk from infection and raise concerns about the importance of the diseases caused by CRKPs [4].

In light of this situation, prevention is one of the most urgent challenges for public health and, although it requires multidisciplinary efforts, early diagnosis of this type of colonisation is crucial to prevent the spread of CRKP and to improve the prognosis of patients [2, 11, 12]. To date, the work-flow for OXA-48 like carbapenemases detection consisted of phenotypic screening, with very low sensitivity for OXA-48 detection due to its activity (PNT-MRN-03 in ISBN-978-84-615-1530-1), followed by a phenotypic or molecular confirmation [2, 8, 13]. However, detection of carbapenem resistance is difficult due to the fact that the presence of carbapenemase genes does not necessarily involve resistance, since their expression is extremely variable. Moreover, some carbapenemase enzymes, such as the OXA-48 like type, show quite low hydrolytic activity, which increases the number of cases of false negative results [11].

Currently, isothermal amplification reactions in association with other techniques that increase the diagnostic specificity, such as clustered regularly interspaced short palindromic repeats (CRISPR)-associated protein (CRISPR-Cas) systems, are becoming more important in the detection of antimicrobial resistant microorganisms [14, 15]. Although different methods of isothermal amplification are available, loop-mediated isothermal amplification (LAMP) reactions display greater specificity than other methods [16, 17] and have previously been used to detect several microorganisms [16, 18-20].

Among the wide variety of endonucleases in CRISPR-Cas systems, the Cas13 enzyme offers several advantages for the development of diagnostic techniques. On the one hand, Cas13 is highly specific and is capable of discriminating between two sequences varying in a single nucleotide [21]. On the other hand, this endonuclease acts on single-stranded RNA (ssRNA) molecules and could be used to detect active infections. The enzyme also shows collateral activity when it is specifically activated, degrading all ssRNA molecules [14, 15]. These features indicate the CRISPR-Cas13 system as a promising tool in the development of diagnostic tests.

In a recent study, our group developed an RT-LAMP-CRISPR-Cas13a technique for the detection of SARS-CoV-2 in nasopharyngeal samples after a process in which no RNA extraction is needed, with the results revealed in HibriDetect strips and yielding 83 % sensitivity and 100 % specificity [22]. Here, we applied this technology to clinical samples from different origins, leading to the successful differentiation of *K. pneumoniae* from other species of the genus Klebsiella, as well as detection of the carbapenem-resistance gene *bla*_OXA*-*48_ (Figure 1).

**Figure 1.**
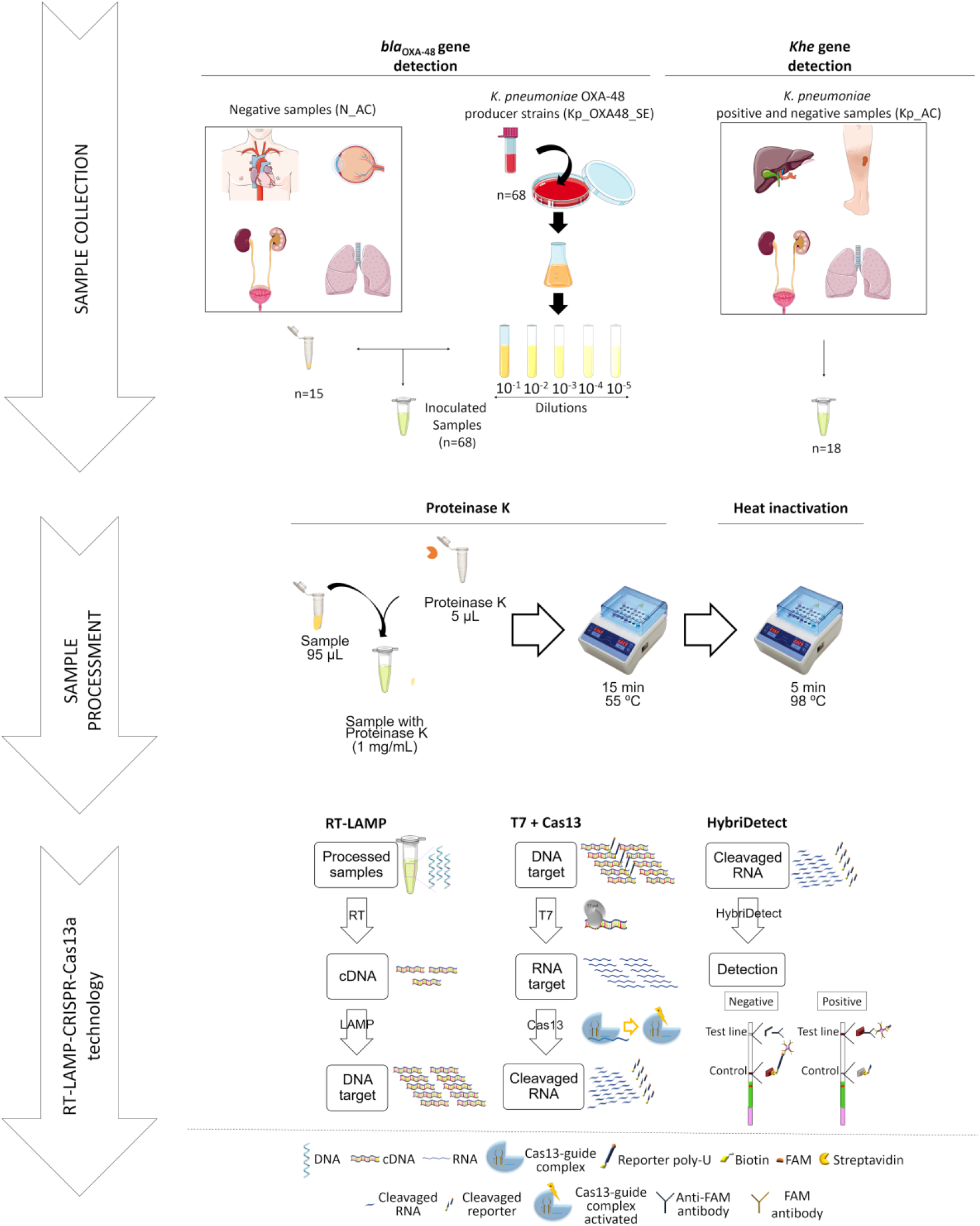
Workflow of the protocol for diagnosis of infectious diseases based on the CRISPR-Cas13a system.

## Material and methods

### Study of the state of the art

A study of the state of the art was conducted with the aim of comparing the use of different classical carbapenemase OXA-48 diagnostic techniques with the novel RT-LAMP-CRISPR-Cas13a technique. First, we compiled a variety of studies of genotypic technologies for *bla*_OXA-48_ gene detection: colorimetric LAMP [23, 24], multiplex PCR [25-27], real time PCR (qPCR) [28], multiplex real time PCR (multiplex qPCR) [29, 30] and multiplex microarray [31]. We then determined the sensitivity, specificity, positive predictive value (PPV) and negative predictive value (NPV) of the different techniques. We used the results to calculate the parameters needed for comparative purposes.

### RT-LAMP primers, crRNAs and RNA reporter

The *K. pneumoniae* house-keeping hemolysin gene (*khe* gene; ID: KX842080.1) and the *bla*_OXA*-*48_ gene (ID: KT175899.1) were selected for the study. The target sequences were analyzed *in silico* with the aim of designing specific primers for the amplification of a conserved genetic region (*khe* gene region 208-396 pb and *bla*_OXA-48_ gene region 174-404 pb). Three pairs of LAMP primers were designed using the PrimerExplorer V5 software (F3/B3, FIP/BIP, Floop/Bloop) to amplify the target regions of the *K. pneumoniae* genome. The FIP LAMP primers included the T7 polymerase promotor in their sequences for the subsequent transcription step. In addition, another pair of PCR primers was designed for each gene (*khe* gene region 190-319 pb and *bla*_OXA-48_ gene region 309-444 pb), and the T7 polymerase promotor was included in the 5’ extreme of the forward primers. Finally, crRNA molecules that hybridize regions of the LAMP and PCR amplicons of both genes were designed (Table 1).

**Table 1.**
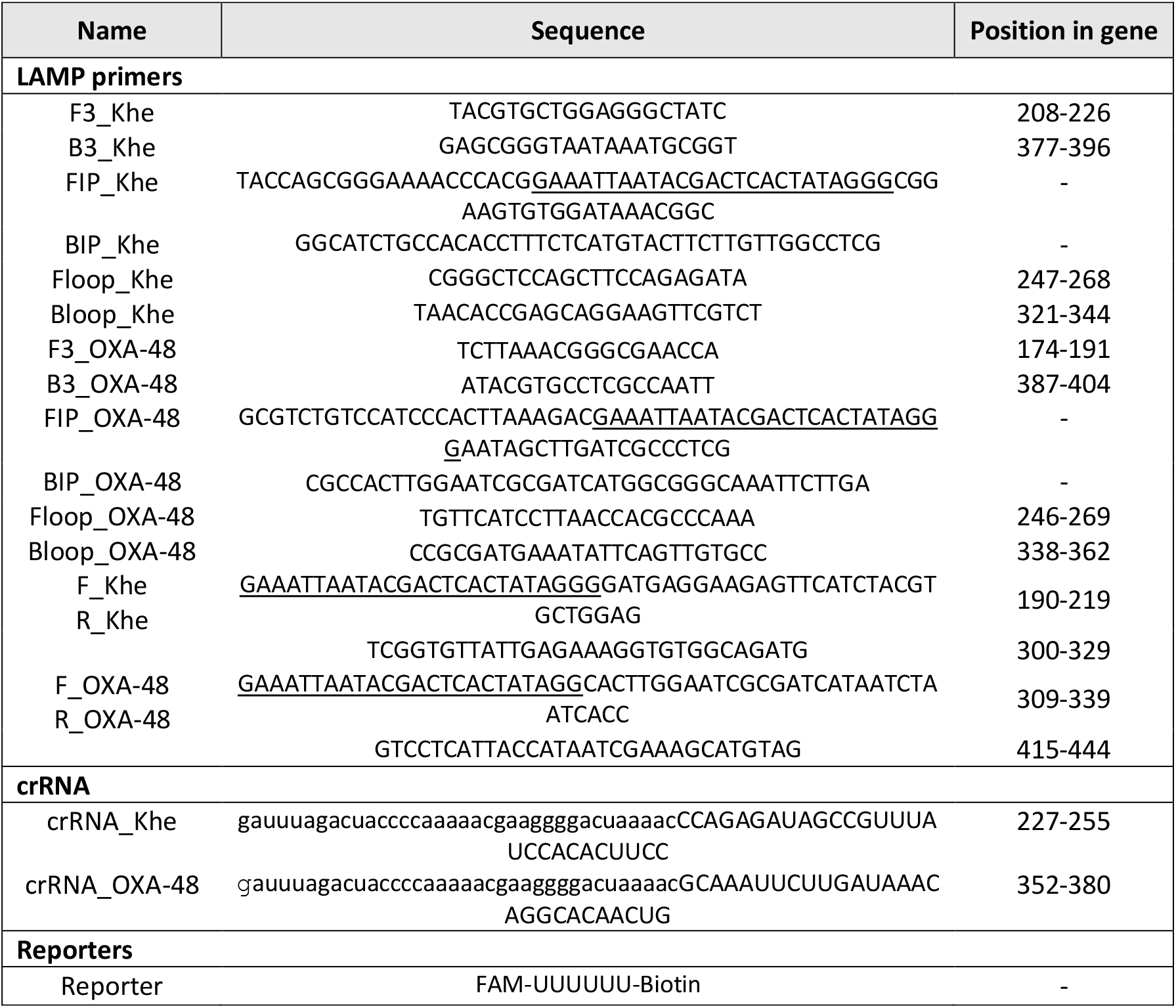
Sequences of primers, crRNAs and reporters. Underlined letters indicate overhang T7 promotor sequence and lower-case letters indicate scaffold sequence. All primers were supplied by IDT and reporters were supplied by GenScript.

The RNA reporter molecules used for signal amplification contained a single isomer derivative of fluorescein modification (FAM) at the 5’
sextreme and a biotin molecule at the 3’extreme (Table 1).

### Clinical samples

The clinical samples used in the study belonged to 2 different groups: (i) samples for *K. pneumoniae* identification (Kp_AC samples, n=18) supplied by the Microbiology Service of the Teresa Herrera Materno Infantil Hospital (A Coruña, Spain). Of these samples, 11 were from patients infected by *K. pneumoniae* non-OXA-48 producer strains (Kp_AC.1-Kp_AC.11) and 7 from patients infected by other Enterobacterales (Kp_AC.12-Kp_AC.18) (Table 2); and (ii) samples for detection of *bla*OXA-48 gene (n=68) from patients without carbapenem-resistant bacterial infection (N_AC samples, n=15) supplied by the Microbiology Service of the Teresa Herrera Materno Infantil Hospital (A Coruña, Spain) and inoculated at random with 68 *K. pneumoniae* OXA-48 producer strains (Kp_OXA48_SE strains) supplied by the Reference Laboratory PIRASOA of the Virgen Macarena Hospital (Sevilla, Spain) (Table 3). In addition, the *K. pneumoniae* ST846-OXA48 strain was used to estimate the minimum quantity (ng) of RNA detectable by the collateral-based detection reaction.

**Table 2.** Kp_AC samples for *K. pneumoniae* identification.

|  | Sample | Strain | Origin |
| --- | --- | --- | --- |
| Positive | Kp_AC.1 | *K. pneumoniae*, *E. coli* and *K. oxytoca* | Urine |
|  | Kp_AC.2 | *K. pneumoniae* | Urine |
|  | Kp_AC.3 | *K. pneumoniae* | Urine |
|  | Kp_AC.4 | *K. pneumoniae* | Urine |
|  | Kp_AC.5 | *K. pneumoniae* | Urine |
|  | Kp_AC.6 | *K. pneumoniae* | Urine |
|  | Kp_AC.7 | *K. pneumoniae* | Bronchial aspirate (BAS) |
|  | Kp_AC.8 | *K. pneumoniae* | Bronchial secretion |
|  | Kp_AC.9 | *K. pneumoniae* | Abscess |
|  | Kp_AC.10 | *K. pneumoniae* | Bronchial biopsy |
|  | Kp_AC.11 | *K. pneumoniae* | Biliary fluid |
| Negative | Kp_AC.12 | *K. aerogenes* | Urine |
|  | Kp_AC.13 | *K. oxytoca* | Sputum |
|  | Kp_AC.14 | *K. oxytoca* | Bronchial secretion |
|  | Kp_AC.15 | *K. aerogenes* | Bronchial secretion |
|  | Kp_AC.16 | *K. oxytoca* | Venous ulcer |
|  | Kp_AC.17 | *K. variicola* | Urine |
|  | Kp_AC.18 | *Raoultella ornithinolytica* | Urine |

**Table 3.**
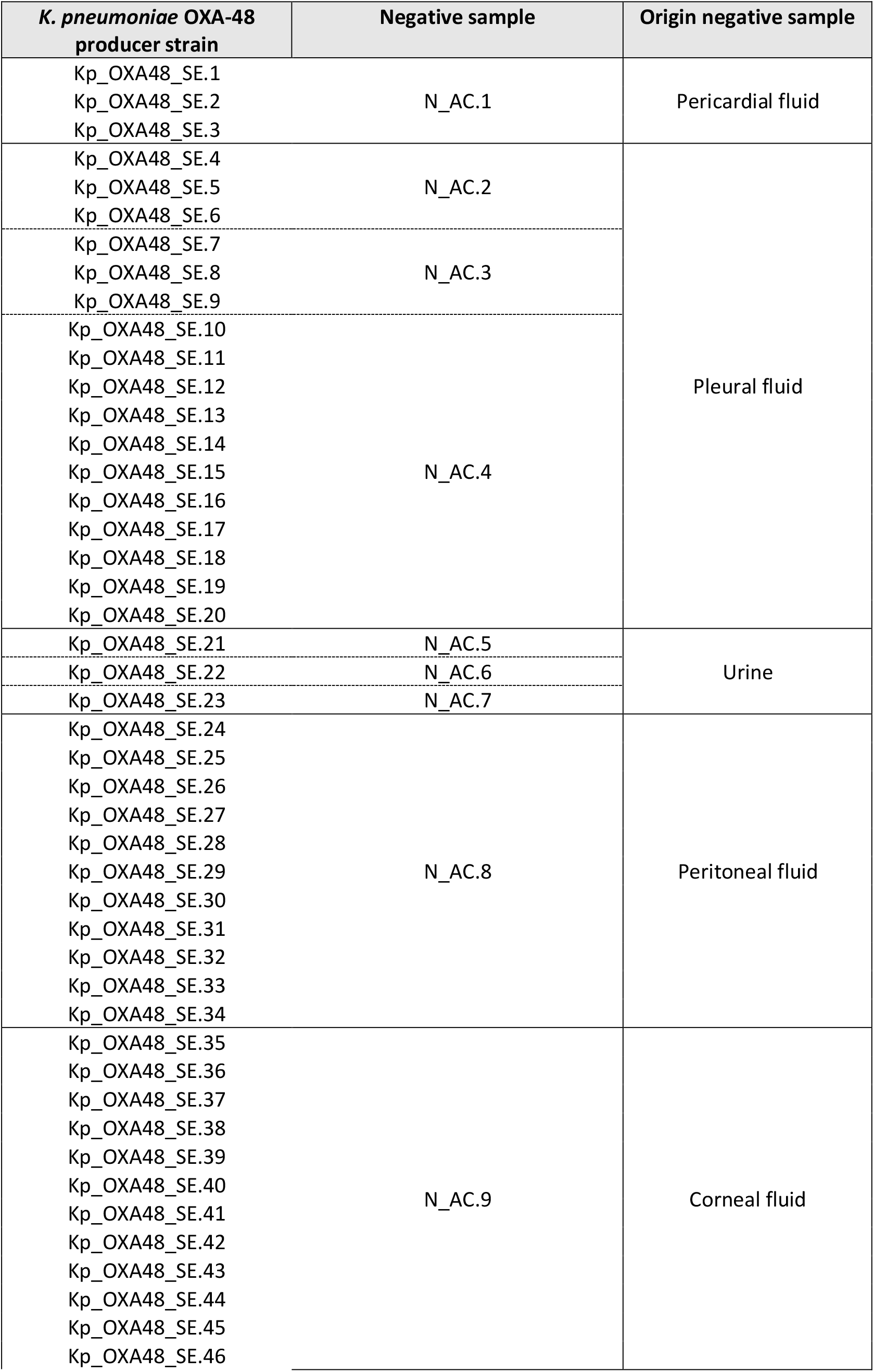
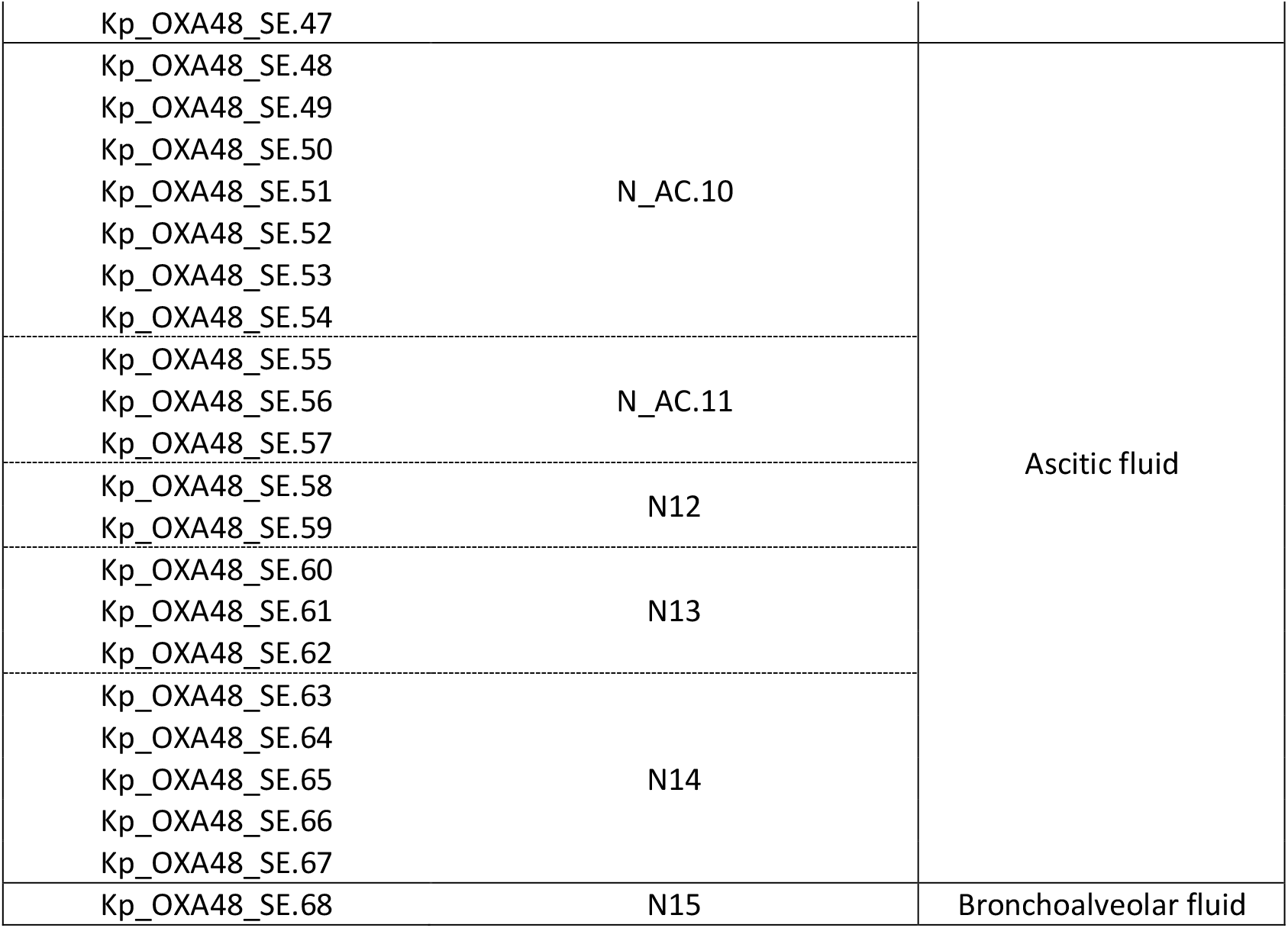
Kp_OXA48_SE strains and N_AC samples for *bla*OXA-48 gene detection through inoculated samples.

### Contamination with *K. pneumoniae* carbapenem-resistant strains

The N_AC samples (n=15) were contaminated at random with 10^4^ CFU/mL of Kp_OXA48_SE strains (n=68). Each N_AC sample was thus inoculated with more than one Kp_OXA48_SE strain (see Table 3). The Kp_OXA48_SE strains were grown in Luria-Bertani (LB) medium (0.5 % NaCl, 0.5 % yeast extract, 1 % tryptone) until an optical density (OD) of 0.6 (10^8^ CFU/mL) was reached, and they were then serially diluted (1:10) in saline solution to yield 10^7^ - 10^5^ CFU/mL. Aliquots (180 µL) of all N_AC samples were used to inoculate 20 µL of dilution 10^−3^ (10^5^ CFU/mL) of the respective Kp_OXA48_SE strains, with the samples to be tested (n=68) finally containing 10^4^ CFU/mL. For example, 3 aliquots were made from the N_AC.1 sample for the inoculation of the Kp_OXA48_SE.1, Kp_OXA48_SE.2 and Kp_OXA48_SE.3 strains, respectively.

### Sample processing

All of the samples, i.e. Kp_AC samples (n=18), N_AC samples (n=15) and the inoculated samples (n=68), were processed following the proteinase K-heat inactivation protocol (PK-HID) in order to obtain the genetic material [32]. Aliquots (95 µL) of the samples were treated for 15 minutes at 55°C with 5 µL of proteinase K (10 mg/mL, Thermo Fisher Scientific, Waltham, MA, USA) prepared at 1 mg/mL in 100 µL of final volume and heat-inactivated at 98 °C for 5 minutes. Finally, processed samples were stored at -80 °C.

### RT-LAMP reaction

DNA was amplified by the RT-LAMP reaction (WarmStart® LAMP Kit (DNA & RNA), NEB, Ipswich, MA, USA) following the manufacturer’ s protocol. Briefly, processed samples (5 µL) were added to a reaction mixture containing 12.5 µL of WarmStart LAMP 2X Master Mix and 2.5 µL of Primer Mix 10X (FIP/BIP 16 µM, F3/B3 2 µM, LOOPF/LOOPB 4 µM, stock) adjusted to a final volume of 25 µL with dH2O. The reactions were incubated at 65 °C for 1 hour.

### Collateral-based detection

Each Cas13a-based detection reaction was incubated at 37 °C for 30 minutes with the following reaction components: 2 µL of cleavage buffer 10X (GenCRISPR™ Cas13a (C2c2) Nuclease, GenScript Biotech Corporation, Piscataway, NJ, USA), 0.5 µL of dNTPs (HiScribe™ T7 Quick High Yield RNA Synthesis Kit, NEB, Ipswich, MA, USA), 0.5 µL of T7 polymerase (HiScribe™ T7 Quick High Yield RNA Synthesis Kit, NEB, Ipswich, MA, USA), 20 U of RNase murine inhibitor (NEB, Ipswich, MA, USA), 0.15 µL Cas13a endonuclease (25 nM, GenCRISPR™ Cas13a (C2c2) Nuclease, GenScript Biotech Corporation, Piscataway, NJ, USA), 0.5 µL crRNA (50 nM, IDT, Coralville, IA, USA), 2 µL of reporter (1000 nM, GenScript Biotech Corporation, Piscataway, NJ, USA) and 5 µL of cDNA sample, adjusted to a final volume of 20 µL with dH_2_O.

### Hybridetect lateral flow

The results were revealed by the HybriDetect lateral flow assay, as described by the manufacturer (Milenia Biotec, Giessen, Germany), with some modifications. Briefly, 20 µL of collateral-based detection product was mixed with 80 µL of assay buffer supplemented with 5 % polyethylenglycol (PEG) in a 96-well plate. Immediately, the gold-extreme of the dipstick was submerged in the mixture and held for 2-3 minutes. Following the manufacturer’ s instructions, reactive strips required calibration before application for management of an optimal RNA reporter concentration.

### Limit of detection (LoD)

In order to estimate the minimum quantity (ng) of RNA target detectable by the collateral-based detection reaction, total DNA was extracted from a *K. pneumoniae* ST846-OXA48 culture (Minutesiprep, Qiagen, Hilden, Germany). The *khe* and *bla*OXA-48 genes were consequently amplified from 5 µL of the extract by PrimeSTAR HS DNA polymerase, following a commercial protocol (Takara Bio, San Jose, CA, USA). Once the amplified regions were purified and quantified by NanoDrop ND-10000 spectrophotometer (NanoDrop Technologies, Waltham, MA, USA), 200 ng of DNA was used for transcription by T7 polymerase (HiScribe™ T7 Quick High Yield RNA Synthesis Kit, NEB, Ipswich, MA, USA). Finally, RNA fragments were purified and quantified by NanoDrop and a collateral-based detection reaction was tested with an RNA gradient in the detection reaction: 1000, 100, 10, 1 and 0.1 ng from each target.

In order to determine the LoD of the CFU/mL inoculated, 3 Kp_OXA48_SE strains (Kp_OXA48_SE.21, Kp_OXA48_SE.36 and Kp_OXA48_SE.37) were grown in LB, until an optical density (OD) of 0.6 (10^8^ CFU/mL) was reached, and they were then serially diluted (1:10) in saline solution to yield 10^8^ to 10^2^ CFU/mL for final inoculation of 20 µL from dilution 10^−2^ (10^6^ CFU/mL) to dilution 10^−5^ (10^3^ CFU/mL) in 180 µL of N_AC samples (N_AC.5 and N_AC.9). The contaminated samples were then tested using the CRISPR-Cas technique.

### *K. pneumoniae* identification by *khe* gene detection

The Kp_AC.1-Kp_AC.11 samples were used to test the capacity of the RT-LAMP-CRISPR-Cas13a technique to identify *K. pneumoniae* by *khe* gene detection, while Kp_AC.12-Kp_AC.18 samples were used as negative controls.

### Carbapenemase OXA-48 identification by *bla*_OXA-48_ gene detection

The 68 inoculated samples were used to test the ability of the RT-LAMP-CRISPR-Cas13a technique to identify the carbapenemase OXA-48 by the *bla*_OXA-48_ gene detection, while the 15 uninoculated N_AC samples were used as negative controls. In both tests, the results obtained, i.e. true positive (TP), false positive (FP), false negative (FN) and true negative (TN), were used to calculate the following parameters: sensitivity (TP/TP+FN), specificity (TN/TN+FP), PPV (TP/TP+FP) and NPV (TN/TN+FN).

## Results

### Study of the state of the art

The best results were obtained in studies that applied colorimetric LAMP technology for *bla*_OXA-48_ and detection of other resistance genes, showing sensitivity, specificity, PPV and NPV values of 100 % in 15 minutes with a fluorescent probe that bound double-stranded DNA (dsDNA) [23] and in 1 hour with a metal ion indicator that changed colour as a consequence of success of the LAMP reaction [24]. The qPCR studies yielded excellent results, with parameter values of 100 %, although they were slower (1-3 hours) [28-30]. On the other hand, the greatest variety of data was obtained in multiplex PCR research, yielding sensitivity values of 92-100 %, specificity values of 88-100 %, PPV values of 21-100 % and NPV of 95-100 %, and taking up to 5 hours [25-27]. Finally, multiplex microarray-based technology yielded sensitivity of 98.3 %, specificity of 99.6 %, PPV of 98.3 % and NPV of 99.6 % and took less time than PCR (2 hours) [31].

### Limit of detection (LoD)

The LoD determination of the collateral-based detection reaction revealed that the CRISPR-Cas13a system detects as few as 1 ng RNA of both *khe* and *bla*_OXA-48_ genes (Figure 2). Furthermore, the LoD was calculated to determine whether the number of CFU/mL of inoculated Kp_OXA48_SE strains was detectable by the RT-LAMP-CRISPR-Cas13a assay. Inoculation of 10^3^ CFU/mL of the strains proved sufficient for detection, although we decided to add 10^4^ CFU/mL to produce a more reliable signal and for better interpretation of the results (Figure 3).

**Figure 2.**
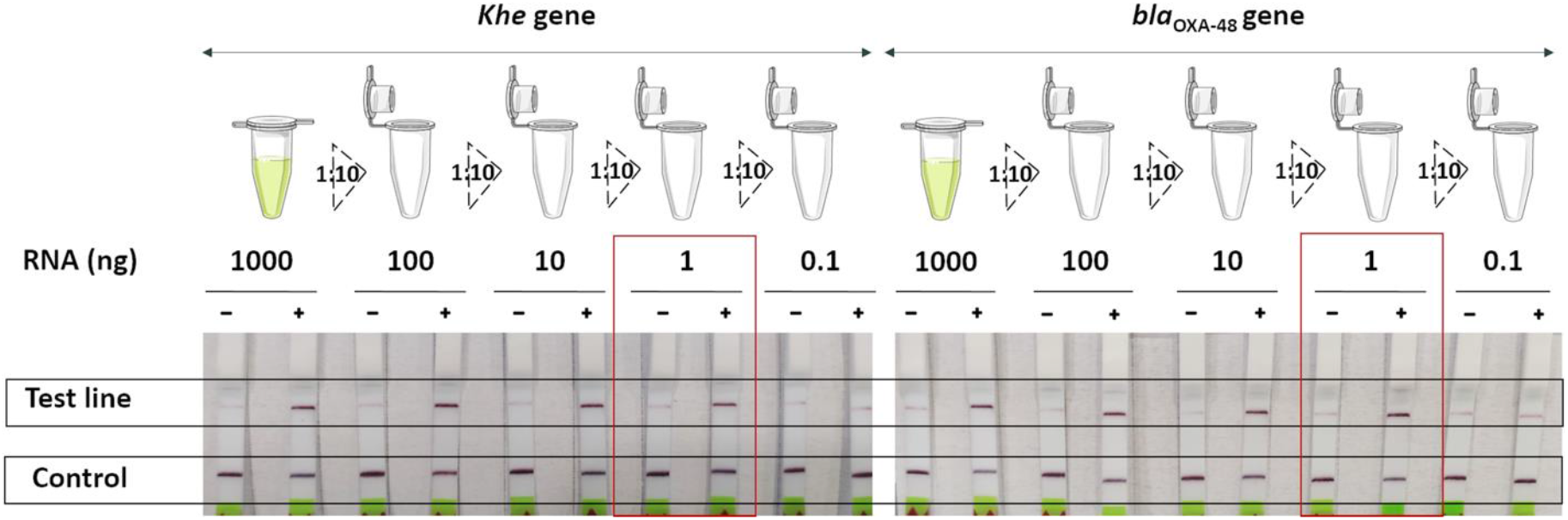
LoD assay for the determination of the quantity (ng) of target that the collateral-based detection reaction identifies using serial dilutions (1:10) from RNA molecules of both *khe* and *bla*OXA-48 PCR amplicons.

**Figure 3.**
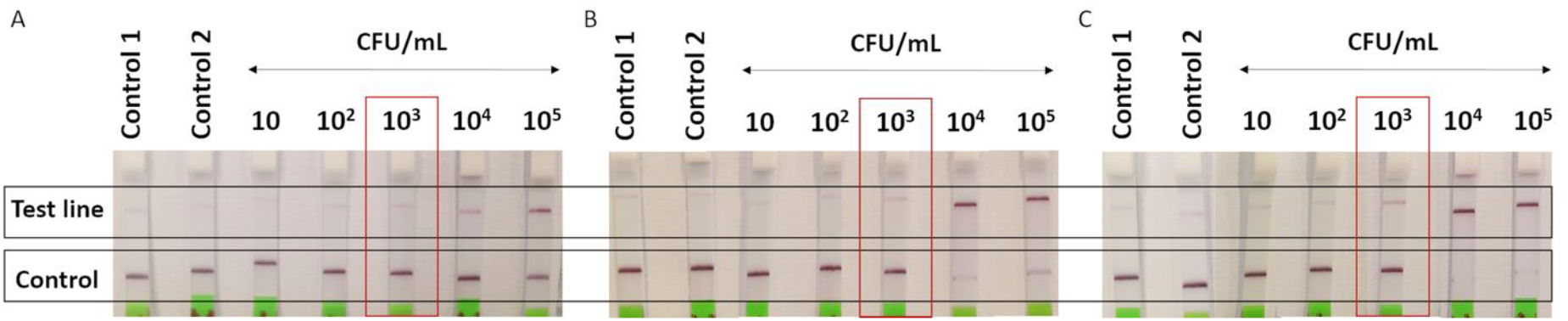
LoD assay for the determination of the CFU/mL of the Kp_OXA48_SE strains inoculated in the N_AC samples that the RT-LAMP-CRISPR-Cas13a technique detects in serial dilutions (1:10) of LB cultures of 3 Kp_OXA48_SE strains: **A**. Kp_OXA48_SE.21 inoculated in the urine sample N_AC.5; **B**. Kp_OXA48_SE.36 inoculated in the cornea sample N_AC.9; **C**. Kp_OXA48_SE.37 inoculated in the urine sample N_AC.9.

### *K. pneumoniae* identification by *khe* gene detection

The RT-LAMP-CRISPR-Cas13a technique was capable of detecting *K. pneumoniae*, even in Kp_AC samples in which it was present with other bacteria. In addition, this technology differentiated *K. pneumoniae* infections from colonization caused by other pathogens, as the results were negative for negative Kp_AC samples (Kp_AC.12 to Kp_AC.18) (Figure 4A). On the basis of the results obtained (Figure 4B), it was estimated that the CRISPR-Cas13a method exhibited 100 % of specificity, sensitivity, PPV and NPV for *K. pneumoniae* identification by the *khe* gene detection (Figure 4C).

**Figure 4.**
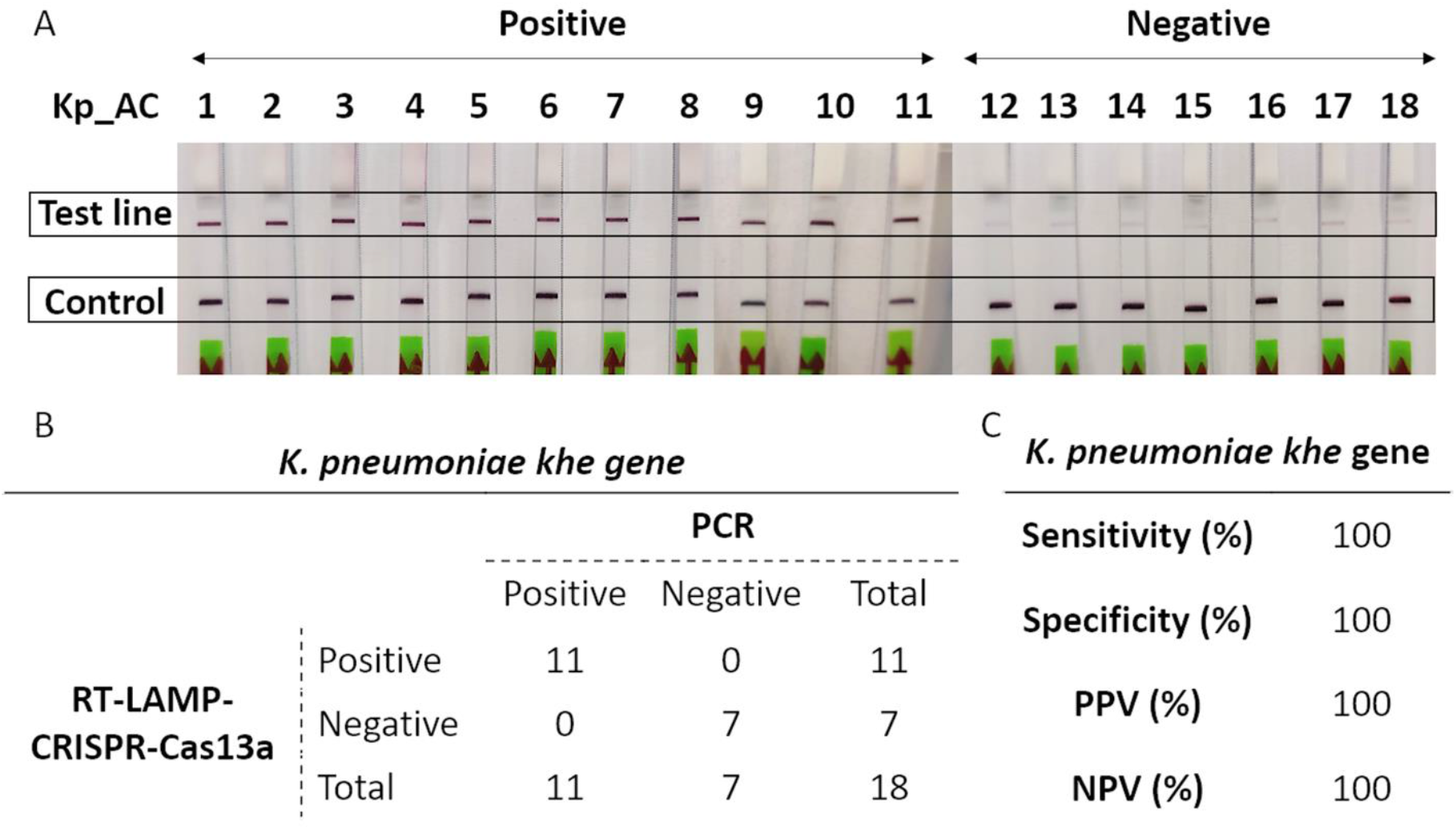
A. Strips for *K. pneumoniae* identification by *khe* gene detection using Kp_AC samples; **B**. Results obtained from *khe* gene detection; **C**. Table including the specificity, sensitivity, PPV and NPV for the RT-LAMP-CRISPR-Cas13a technique from processing of data in Figure 4B.

### Carbapenemase OXA-48 identification by *bla*OXA-48 gene detection

Negative results were obtained by analysis of the 15 N_AC samples by the RT-LAMP-CRISPR-Cas13a technique. The technique also correctly detected the *bla*OXA-48 gene in the inoculated samples (Figure 5A). Based on the results obtained (Figure 5B), we estimated that the LAMP-CRISPR-Cas13a method exhibits 100 % of specificity, sensitivity, PPV and NPV for *bla*OXA-48 gene detection (Figure 5C).

**Figure 5.**
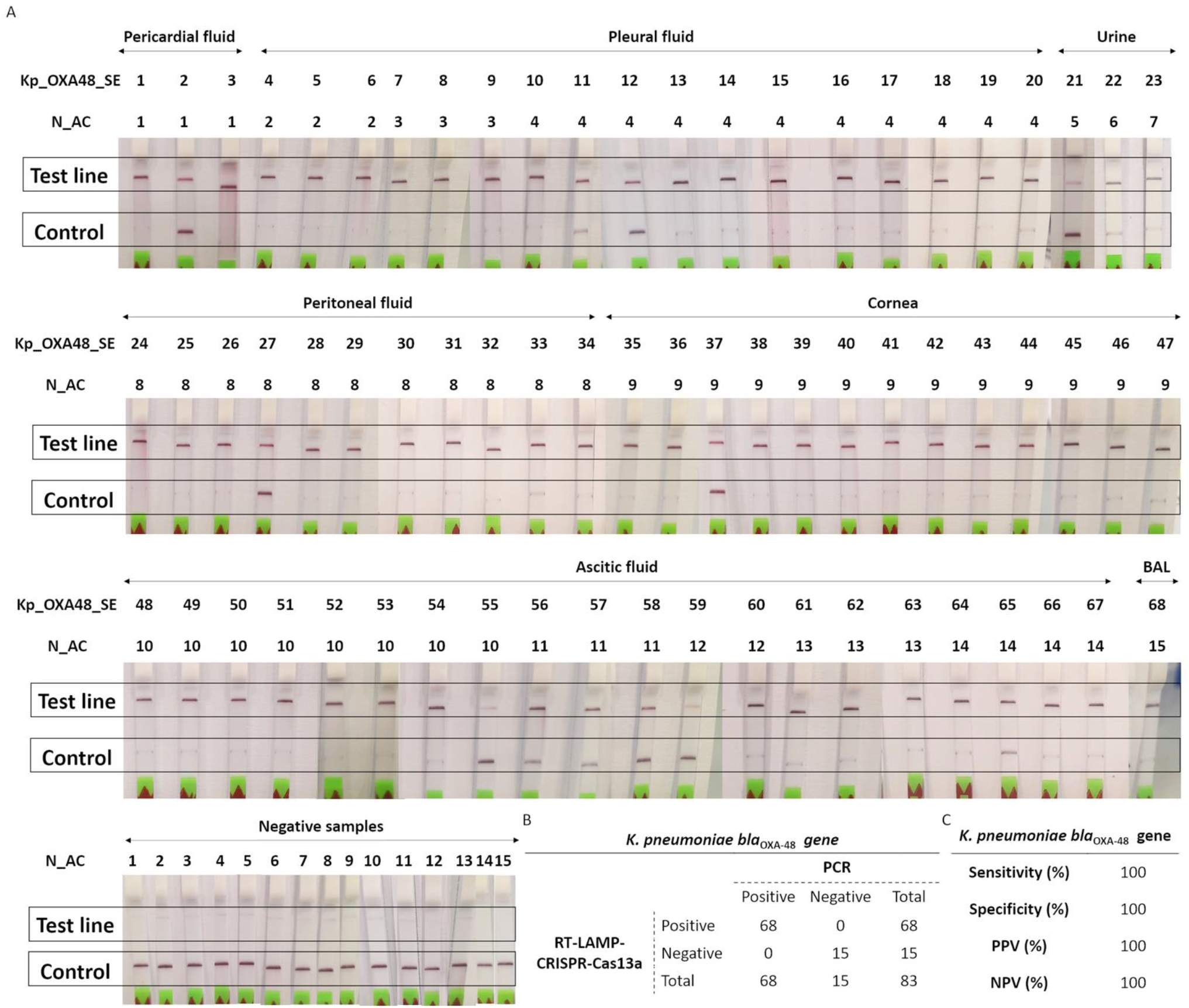
A. Strips for carbapenemase OXA-48 identification by *bla*OXA-48 gene detection using inoculated samples as positive samples and N_AC samples as negative controls; **B**. Results obtained from *bla*OXA-48 gene detection; **C**. Table including the specificity, sensitivity, PPV and NPV for the RT-LAMP-CRISPR-Cas13a technique from processing data in Figure 5B.

## Discussion

Study of the state of the art revealed that greater efforts must be made to innovate in genotypic diagnostic methods for carbapenem-resistance, due to the important repercussions that this has in public health and to the current lack of a technique that combines high levels of sensitivity and specificity while reducing the associated costs and time (Table 4). As stated above, the presence of a carbapenemase gene does not involve resistance, as the level of expression in these strains is highly variable [11]. Therefore, the development of a diagnostic technique that detects target RNA molecules, as in the RT-LAMP-CRISPR-Cas13a technique, will enable the identification of strains that actually produce the carbapenemase enzymes, irrespective of the presence of the carbapenemase genes.

**Table 4.** Percentages of sensitivity, specificity, PPV and NPV parameters reported in 9 articles on the application of genotypic methods for carbapenemase genes detection.

| Technique | Time | Sample/<br>Isolate | Extraction<br>kit | Sensitivity<br>(%) | Specificity<br>(%) | PPV<br>(%) | NPV<br>(%) | Ref |
| --- | --- | --- | --- | --- | --- | --- | --- | --- |
| LAMP<br>fluorescent | 15 min | Isolate | No | 100 | 100 | 100 | 100 | [23] |
| LAMP –HNB | 1 h | Isolate | No | 100 | 100 | 100 | 100 | [24] |
| Multiplex PCR | <1.5 h | Sample | No | 100 | 88 | 21 | 100 | [25] |
| Multiplex PCR | 3 h | Sample | Yes | 92.5 | 100 | 100 | 95.2 | [26] |
| Multiplex PCR | 5 h | Isolate | Yes | 100 | 99.4 | 93.9 | 100 | [27] |
| qPCR | 3 h | Isolate | No | 100 | 100 | 100 | 100 | [28] |
| Multiplex qPCR | <1 h | Isolate | Yes | 100 | 100 | 100 | 100 | [29] |
| Multiplex qPCR | <1.5 h | Isolate | No | 100 | 100 | 100 | 100 | [30] |
| Multiplex<br>microarray | 2 h | Sample | No | 98.3 | 99.6 | 98.3 | 99.6 | [31] |

In this work, our research group has developed an RT-LAMP-CRISPR-Cas13a protocol for diagnosing *K. pneumoniae* OXA-48 producers infected with a LoD of 10^3^ CFU/mL (Figure 3). Although in a previous study the LoD of the LAMP technique for *K. pneumoniae* identification was 10^2^ CFU/mL [33], this difference can be explained by the fact that the CRISPR-Cas systems add a level of specificity to the diagnostic tests that the LAMP reaction alone does not have. On the other hand, another assay yielded a LoD of 10^4^ CFU/mL for *bla*_OXA-48_ gene detection by the LAMP reaction, while the LoD was 10^9^ CFU/mL for PCR detection [24]. In addition, detection by the CRISPR-Cas13a system yielded a LoD of 1 ng (Figure 2), which may indicate that the diagnostic protocol could be directly applied to samples without prior amplification by RT-LAMP, thus reducing the time for diagnosis and the cost of material required.

Considering the criteria recommended by the WHO (UNICEF/UNDP/World Bank/WHO Special Programme for Research and Training in Tropical Diseases & World Health Organization. (2010). Accessible quality-assured diagnostics: annual report 2009. World Health Organization. https://apps.who.int/iris/handle/10665/84529), the novel technique fulfils the three key features of accuracy, accessibility and affordability. This is because, on the one hand, it showed an accuracy ((TP+TN)/total) of 100 % for *K. pneumoniae* identification and *bla*_OXA-48_ gene detection, respectively. Furthermore, it is both accessible and affordable as neither specific equipment nor trained personnel are required, and the amounts of enzymes needed per reaction are quite low.

Comparing our results on sensitivity, specificity, PPV and NPV with those obtained in previous studies, we found that the specificity and PPV values of the RT-LAMP-CRISPR-Cas13a technique were higher than those in 2 out of 3 multiplex PCR studies reviewed [25, 27], and the sensitivity and NPV values were also higher than in 1 of 3 of these studies [26]. Moreover, this technique showed higher sensitivity, specificity, PPV and NPV values than those from multiplex microarray study reviewed [31]. Among the others, the RT-LAMP-CRISPR-Cas13a protocol produced results as good as those obtained with LAMP and qPCR assays [23, 24, 28-30]. However, both LAMP studies used probes that gave a confirmatory signal of LAMP amplification without the corroboration of the sequence-specificity of the amplicon, which is worrisome due to the possible non-specificity of the isothermal amplification reaction [23, 24]. However, CRISPR-Cas based technologies use specific probes for the different targets, contributing to the development of a highly specific diagnostic test. Additionally, one multiplex qPCR study used a SYBR green probe [29], which bound any dsDNA molecules, and its specificity was therefore extremely low when applied to clinical samples. Moreover, SYBR green probes inhibit LAMP reaction and must be added after the end of the reaction, increasing the risk of contamination [24].

In summary, the high levels of specificity, sensitivity, PPV and NPV obtained with this promising RNA-extraction free protocol makes the LAMP-CRISPR-Cas13a system one of the best rapid and specific diagnostic methods for infectious diseases. Thus, this technique could be established as a diagnostic tool in the routine clinical detection of other infectious diseases, such as those caused by multiresistant pathogens. However, Cas13 detection methods should be optimized to enable direct diagnosis without prior amplification of nucleic acids.

## Acknowledgment

This study was funded by grant PI19/00878 awarded to M. Tomás, within the State Plan for R+D+I 2013-2016 (National Plan for Scientific Research, Technological Development and Innovation 2008-2011) and co-financed by the ISCIII-Deputy General Directorate for Evaluation and Promotion of Research - European Regional Development Fund “A way of Making Europe” and Instituto de Salud Carlos III FEDER, Spanish Network for the Research in Infectious Diseases (REIPI, RD16/0016/0006 and CIBER CB21/13/00012, CB21/13/00084 and CB21/13/00095), Instituto de Salud Carlos III FEDER Experiments were funded by grant IN607D 2021/10 within the GAIN (Agencia Gallega de Innovación) and by the Study Group on Mechanisms of Action and Resistance to Antimicrobials, GEMARA (SEIMC, http://www.seimc.org/) and finally, ESCMID grant (European Society of Clinical Microbiology and Infectious Diseases) awarded to L. Fernández-García. O. Pacios, L. Fernández-García and M. López were financially supported by the grants IN606A-2020/035, IN606B-2021/013 and IN606C-2022/002, respectively (GAIN, Xunta de Galicia). I. Bleriot was financially supported by pFIS pro-511gram (ISCIII, FI20/00302).

